# PleistoDist: A toolbox for visualising and quantifying the effects of Pleistocene sea-level change on island archipelagos

**DOI:** 10.1101/2022.05.13.491891

**Authors:** David J.X. Tan, Ethan F. Gyllenhaal, Michael J. Andersen

**Affiliations:** Department of Biology and Museum of Southwestern Biology, University of New Mexico, Albuquerque, USA

**Keywords:** Bathymetry, Biogeography, Land bridges, Islands, Pleistocene, Sea level, Phylogeography, Gene flow

## Abstract

1. Pleistocene sea-level change played a significant role in the evolution and assembly of island biotas. The formation of land bridges between islands during Quaternary glacial maxima, when sea levels were up to 120 metres below present-day sea levels, often facilitated historical dispersal and gene flow between islands that are today geographically disconnected.
2. Despite this, relatively few studies have attempted to quantify the effects of Pleistocene sea-level change on the evolution of island species assemblages.
3. Here we present PleistoDist, an R package that allows users to visualise and quantify the effects of Pleistocene sea-level change on islands over time, and test multiple temporally explicit hypotheses of inter-island dispersal and community assembly.
4. Re-analysing published datasets, we demonstrate how using PleistoDist to account for historical sea-level change can provide greater explanatory power when analysing extant island communities, and show how population genetic simulations can be used to generate spatiotemporally explicit neutral expectations of population genetic structure across island archipelagos.

## 1. Introduction

One of the key insights of the theory of island biogeography (MacArthur and Wilson 1967) is that the size and orientation of islands play a significant role in structuring the evolution and assembly of island communities. Specifically, the theory predicts that proximate islands experience higher inter-island immigration, and consequently increased gene flow and community similarity (Kimura and Weiss 1964; Simberloff 1974), and that larger islands tend to have more species (Gleason 1922). Yet, empirical species distributions across island archipelagos highlight numerous exceptions. At macroecological scales, the distributions of terrestrial megafauna across the Sundaic and Wallacean regions (Wallace 1869; Lohman et al. 2011) and the decoupling of species-area correlations on land-bridge islands in the Aegean Sea (Hammoud et al. 2021) illustrate how land bridges can facilitate dispersal, colonisation, and biotic homogenisation between distant islands. Phylogeographic studies have also demonstrated that patterns of intraspecific genetic differentiation within island archipelagos are correlated with land-bridge connectivity (Leonard et al. 2015; Cros et al. 2020).

Incorporating historical effects of Pleistocene sea-level change into models of island biogeography is thus essential to better understanding the patterns of community assembly and gene flow observed in island archipelagos (Lohman et al. 2011). However, while many studies have qualitatively described the impact of transient land bridges in island systems (Reilly et al. 2019; Cros et al. 2020), few have addressed this issue quantitatively (see Darwell et al (2020); Sin et al (2022) for notable exceptions). To address the need for methods aimed at quantifying the effects of sea-level change on islands, we present PleistoDist (https://github.com/g33k5p34k/PleistoDistR), an R package that allows users to visualise sea-level change over time and calculate multiple metrics of island shape, inter-island distance, net inter-island migration, and inter-island visibility, all normalised over Pleistocene time. This tool leverages the availability of high-resolution bathymetric data sources and generates outputs that can be used for a wide variety of downstream analyses.

## 2. PleistoDist Workflow

### 2.1 Input Files

PleistoDist requires two main input files: a bathymetry raster to generate maps of island extents at different historical sea levels and a shapefile of source points from which to calculate island metrics and/or pairwise distances. By default, users are advised to use bathymetry data from the General Bathymetric Chart of the Oceans (GEBCO: https://www.gebco.net), a global database of Earth’s terrestrial and undersea terrain at 15-arc-second resolution, although PleistoDist can accept any type of ASCII-formatted bathymetry data as input. Users must also specify a map projection appropriate to their area of interest, as well as a time cutoff to set the temporal scope of the analysis.

### 2.2 Generating the interval file

PleistoDist models historical sea-level change by decomposing Pleistocene sea levels (by default from Bintanja and van de Wal (2008)) into a number of discrete intervals for a user-defined period of time. These intervals can be calculated in two different ways: by binning over time with the *getintervals_time* function, or by binning over sea level with the *getintervals_sealvl* function (Fig. 1). Binning over time involves dividing the timespan of interest into several equal time intervals and calculating the mean sea level for each interval (Fig. 1A). In contrast, binning over sea level involves calculating the total range of sea-level change for the timespan of interest and dividing this range into several equal bins (Fig 1B). For the latter method, the mean sea level of each bin is the average of the minimum and maximum sea level for that bin, without respect to time. Once calculated, the intervals are written to an ‘interval file’ (Fig. 1) in comma separated value (CSV) format in the output folder.

**Figure 1:**
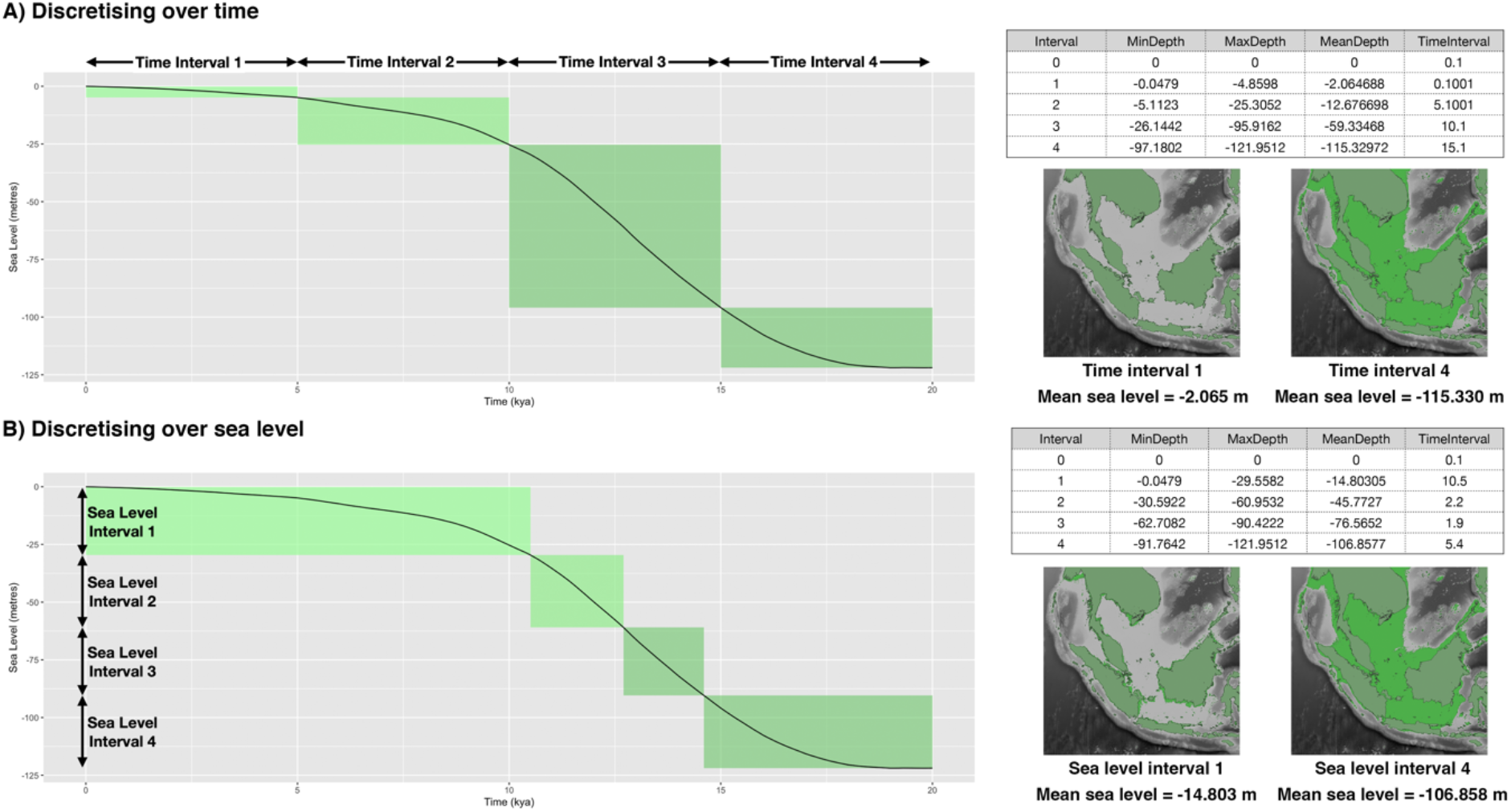
PleistoDist provides two different methods for discretising sea-level change, either by time (A) or by sea level (B), as illustrated here for Sundaland with a time cutoff of 20 kya, five intervals.

### 2.3 Generating maps of island extents

Users can generate maps of island extents based on mean sea levels specified in the interval file (Fig. 1) using the *makemaps* function. This function reprojects the input bathymetry raster into the user-specified map projection using a bilinear resampling method and generates raster and shapefile outputs of island extents in three formats: a shapefile of island polygons, an ASCII flat raster with no topography, and an ASCII topographic raster that preserves the original elevations of each land pixel. Since the *makemaps* function reads intervals and mean sea levels directly from the interval file, users can customise the output maps generated by this module by manually editing the interval file.

### 2.4 Calculating intra-and inter-island metrics

Based on the maps generated, PleistoDist can calculate a variety of intra- and inter-island metrics for each time or sea-level interval, as well as calculate the weighted mean of these values over Pleistocene time. For metrics of island shape, PleistoDist includes functions that calculate island perimeter, area, and surface area for each interval in the interval file. As for inter-island distances, PleistoDist can calculate three kinds of island-to-island distances (centroid-to-centroid, least shore-to-shore, and mean shore-to-shore distances; Fig. 2), as well as two kinds of point-to-point distances (Euclidean and least-cost distances; Fig. 3) for each interval specified in the interval file. Islands that merge at particular intervals are considered equivalent for these and future metrics.

**Figure 2:**
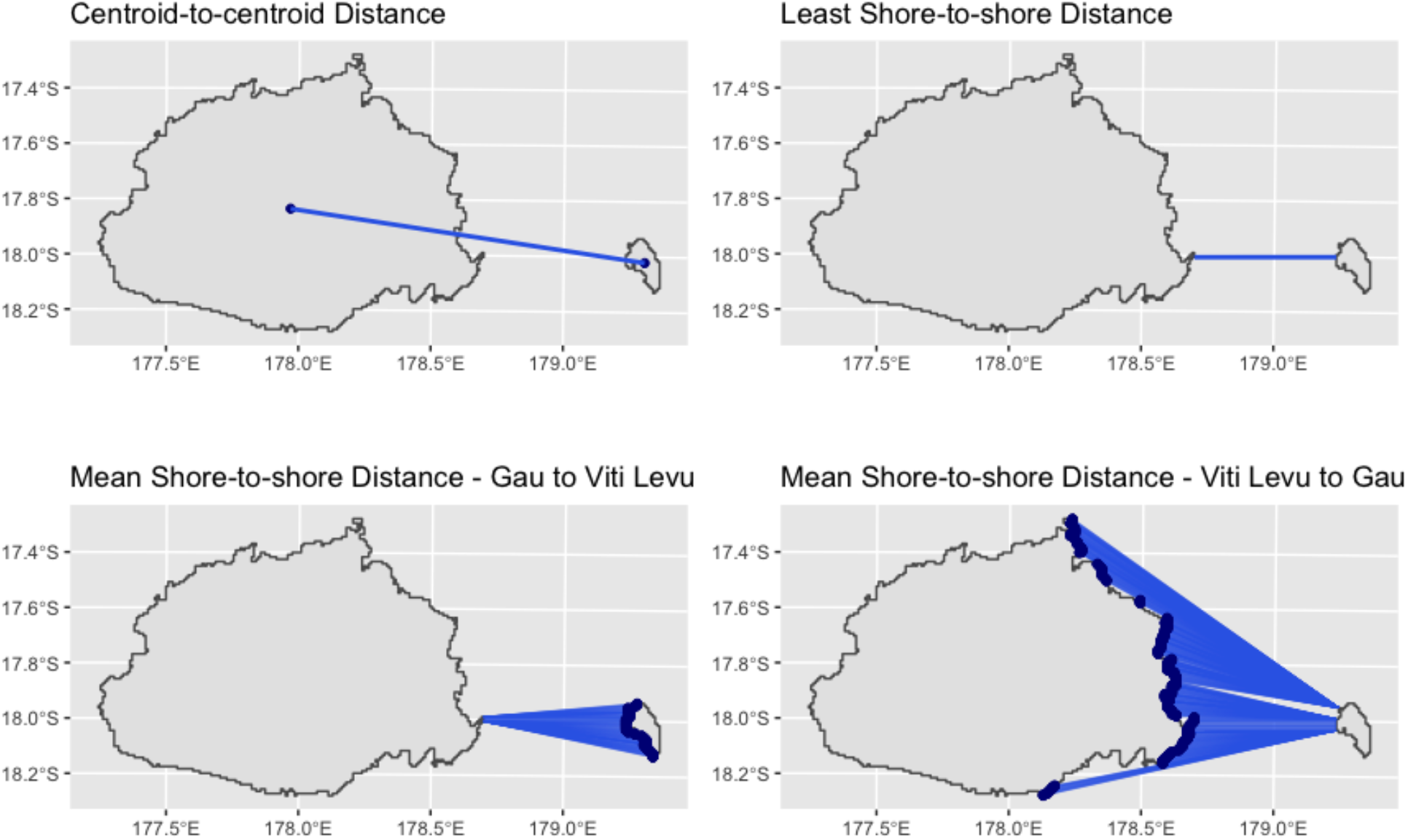
PleistoDist calculates three inter-island distances: centroid-to-centroid distance, least shore-to-shore distance, and mean shore-to-shore distance, illustrated here with Viti Levu (left) and Gau islands, Fiji. Note how inter-island distances are asymmetric for the mean shore-to-shore distance.

**Figure 3:**
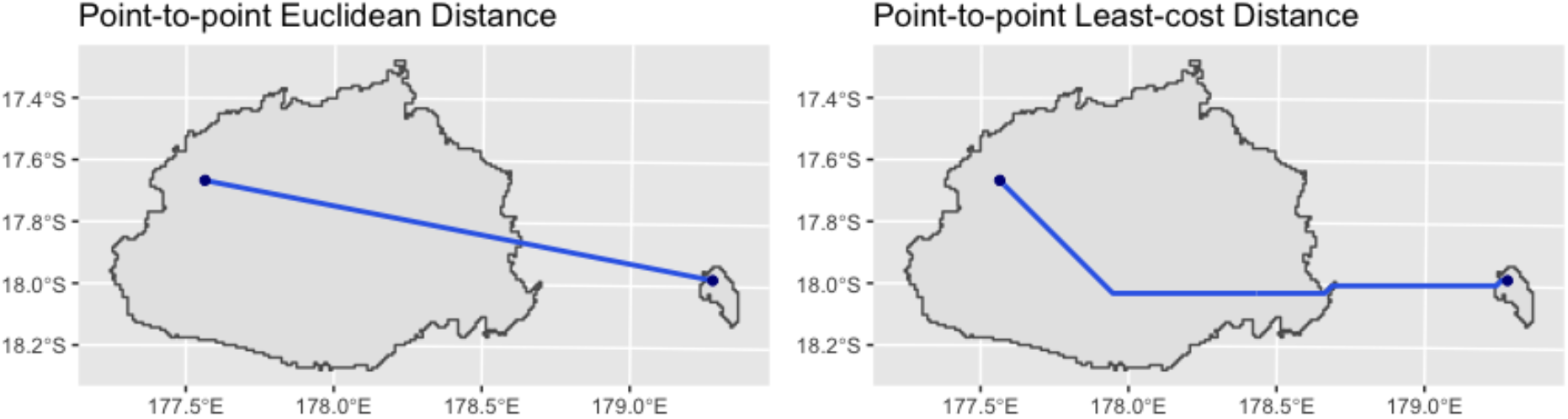
PleistoDist calculates two distance measures between source points: Euclidean distance between points (as the crow flies, invariant across all intervals), and least-cost distance (which minimises overwater movement).

These metrics of inter-island distance allow users to test different models of isolation-by-distance, while accounting for changes in inter-island distance over time.

In addition to metrics of island shape and inter-island distance, PleistoDist includes two higher-level functions that calculate the expected equilibrium net migration between island pairs (MacArthur and Wilson 1967), as well as the visibility of an island relative to an observer on another. The net migration calculation is based on a model described by MacArthur and Wilson (1967) estimating the number of successful propagules dispersing from a source island (island1) to a sink (island2), as given by the equation:

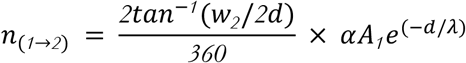

where *n*_(*1*→*2*)_ is the number of propagules of a particular species successfully dispersing from island1 to island2, *w*_*2*_ is the width of island2 relative to island1, *d* is the distance between the two islands, *α* is the population density of the propagule population on island1, *A*_*1*_ is the area of island1, and *λ* is the mean overwater dispersal distance of the propagule (see Gyllenhaal et al. 2020 for further discussion). While this equation is difficult to solve since it requires estimates for both *α* and *λ*, if we take the ratio of *n*_(*1*→*2*)_ and *n*_(*2*→*1*)_—the net migration between islands 1 and 2—we cancel out both the *α* and exponential terms and reduce the relationship to:

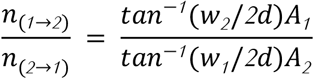

assuming that *d, λ*, and *α* are identical for both islands. Since the parameters of this equation contain only measures of island width, distance, and area, it can easily be solved by PleistoDist-generated outputs. Therfore if 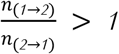, there should be a net movement of individuals from island1 to island2 and vice versa under neutral expectations of the theory of island biogeography.

The inter-island visibility calculation is based on an estimate of the maximum sightline (i.e. horizon distance) of an observer on an island, given by the equation:

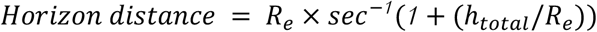

where *h*_*total*_ is the sum of the ground elevation and the height of the observer above the ground, and *R*_*e*_ is the Earth’s radius for that particular inter-island great circle arc, accounting for sea-level change, approximated by the equation:

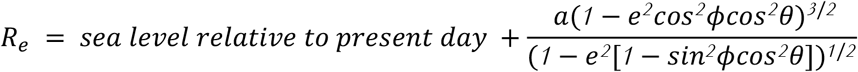

where *a* is Earth’s equatorial radius, *e*^*2*^ is the squared eccentricity of Earth, ϕ is the approximate azimuthal angle between the line connecting the observer and the destination island and Earth’s North-South axis, and θ is the parametric latitude of the observer (Helman 2012). If the destination island falls within the horizon distance radius, PleistoDist performs a viewshed analysis to estimate the observer-visible area of the destination island, accounting for occluding effects of mountains and other geographical features (Fig. 4).

**Figure 4:**
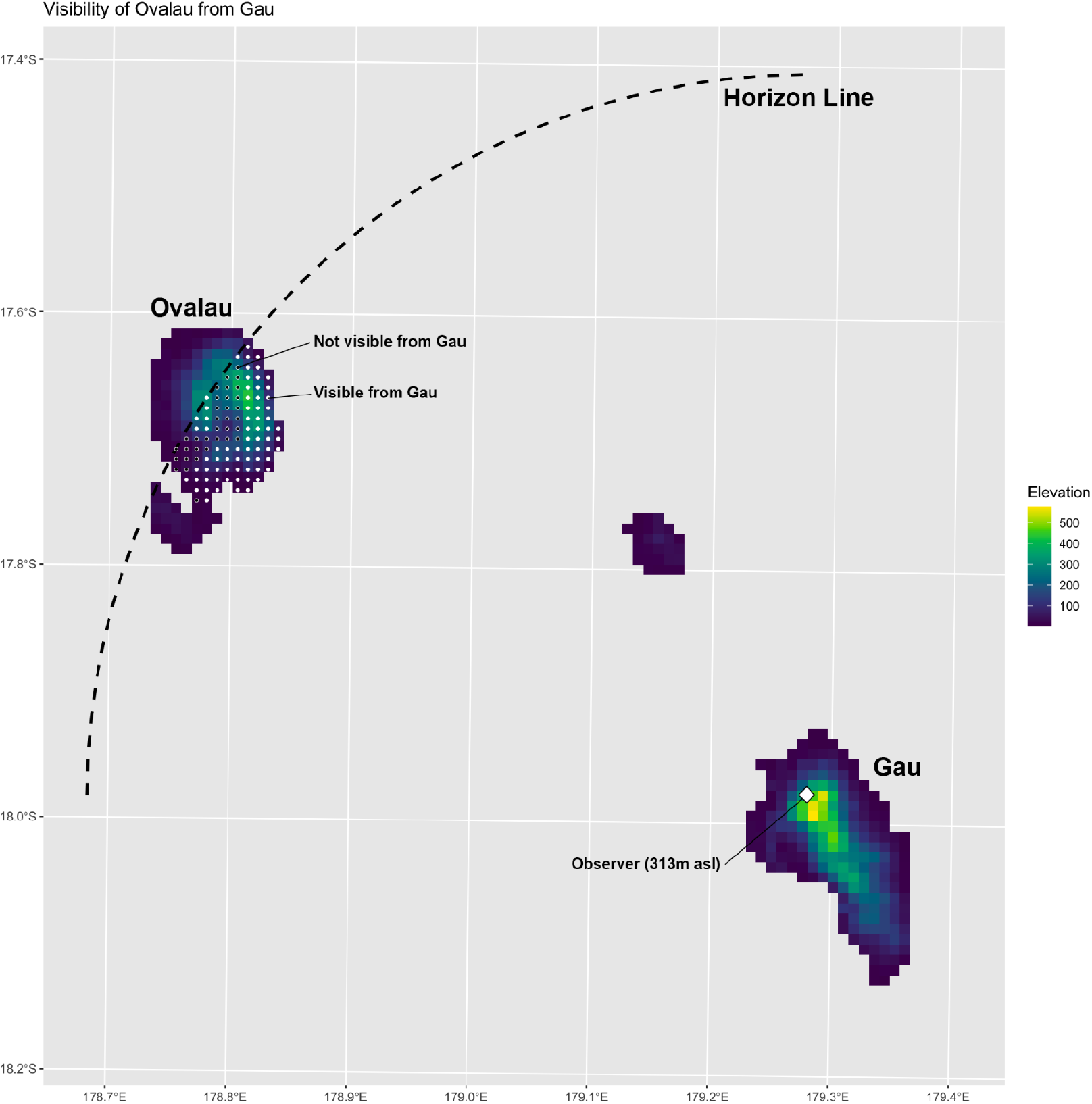
PleistoDist estimates the visibility of a destination island relative to an observer on an origin island by calculating the horizon distance and performing a viewshed analysis to estimate the visible non-occluded area of the destination island.

## 3. PleistoDist Applications and Case Studies

### 3.1 Isolation-by-distance in Caribbean flightless ground crickets (*Amphiacusta sanctaecrucis*)

Papadopoulou and Knowles (2015) studied the effects of Pleistocene island connectivity on the genetic differentiation patterns of *Amphiacusta sanctaecrucis* populations in the Virgin Islands, and found that population divergence times broadly correlate with a period of fluctuating sea levels and inter-island land-bridge connections (∼75–115 kya). In addition, all Virgin Island populations appeared to exhibit a pattern of isolation-by-distance, except those from St. Croix.

Because these analyses were based on present-day Euclidean distances, we used PleistoDist to replicate the landscape genetic analyses using geographical distance matrices that account for sea-level change over time, for a time cutoff of 20 kya (corresponding to the last glacial maximum) and 20 time intervals. Similar to Papadopoulou and Knowles (2015), Mantel tests show no significant correlation between genetic and Euclidean distance (R^2^ = 0.228273, p = 0.0521429; Table 1) when all populations are included in the analysis (Table 1). None of the other geographic distance metrics showed a significant Mantel correlation with genetic distance either. However, given the susceptibility of Mantel tests to spatial autocorrelation (Legendre et al. 2015), it is probable that genetic variation is non-linearly distributed across geographic space. Using distance-based redundancy analyses (dbRDA), which account for spatial autocorrelation, we found that unlike Papadopoulou & Knowles (2015), we observed a highly significant correlation between genetic distance and time-corrected least shore-to-shore and centroid-to-centroid distances (p = 0.00661 and 0.007043, respectively; Table 1), while Euclidean distance shows a marginally significant correlation with genetic distance (p = 0.034779; Table 1). Further model selection indicated that the time-corrected centroid-to-centroid distance best fit the variation observed in the genetic data, suggesting that genetic divergence patterns were likely structured by broad-scale isolation-by-distance driven by inter-island overwater dispersal between panmictic populations.

**Table 1:** Results of Mantel tests and dbRDA for *Amphiacusta sanctaecrucis* populations across the Virgin Islands for different metrics of geographic distance.

| Distance Type | Mantel $R^2$ | Mantel p-value | dbRDA $R^2_{adj}$ | dbRDA p-value |
| --- | --- | --- | --- | --- |
| Euclidean distance | 0.228273 | 0.0521429 | 0.3509484 | 0.034779 |
| Least-cost distance | 0.2203824 | 0.0521409 | 0.1387593 | 0.3025617 |
| Least shore-to-shore distance | 0.0426482 | 0.1997118 | 0.4736347 | 0.00661 |
| Centroid-to-centroid distance | 0.0090014 | 0.2004748 | 0.5933486 | 0.007043 |
| Mean shore-to-shore distance | 0.0035279 | 0.2976187 | -0.4337182 | 0.9366181 |

### 3.2 Species-area relationships of Aegean Island angiosperms

Hammoud et al. (2021) examined the relationship between island area and species richness in the Aegean Islands across multiple angiosperm taxa, taking into account historical sea-level change over the Quaternary. They find that true islands, which were historically unconnected to the mainland, exhibit stronger correlations between island area and endemic species richness relative to land-bridge islands.

Using PleistoDist, we reanalysed angiosperm species-area relationships on the true islands of the Aegean sea using both contemporary and time-corrected estimates of island area for 40 depth intervals over the last 20 kya. We found that species richness is significantly positively correlated with island area for all angiosperm chorotypes (Table 2), consistent with the findings of Hammoud et al. (2021). However, contrary to Hammoud et al. (2021), our extended analyses show that present day 2D island area is the best predictor of angiosperm species richness instead of time-corrected 2D area (Table 2), suggesting that species turnover rates on true islands are fast, resulting in a rapid equilibration of species richness to contemporary island extents.

**Table 2:** Results of simple linear regressions between species richness and island area, for different angiosperm chorotypes and island area metrics.

| Chorotype | Present Day<br>2D Area $R^2_{adj}$ | p-value | AIC | Time-corrected<br>2D Area $R^2_{adj}$ | p-value | AIC |
| --- | --- | --- | --- | --- | --- | --- |
| Total Species<br>Richness | 0.7621572 | 8.51e-17 | 22.043369 | 0.3917022 | 7.04e-7 | 68.9961019 |
| Native Non-<br>endemics | 0.7445056 | 4.80e-16 | 26.3768068 | 0.3873976 | 8.38e-7 | 70.1025982 |
| Endemics | 0.4888763 | 9.80e-9 | 81.6361191 | 0.2662759 | 7.45e-5 | 99.7121938 |

### 3.3 Inter-island migration of *Horornis* bush-warblers across the Fijian archipelago

One of the predictions of the theory of island biogeography (MacArthur & Wilson, 1967) is that the relative rate of migration between island pairs can be predicted by the size, relative orientation, and distance of source islands. Gyllenhaal et al. (2020) tested this expectation by estimating the rates of inter-island migration in the Fiji bush warbler (*Horornis ruficapilla*) between the four large islands of Fiji, concluding that rates of inter-island migration are largely consistent with neutral expectations. This neutral expectation can now be easily and accurately calculated using PleistoDist, while taking into account the effect of sea-level change over Pleistocene time.

Re-running this analysis using time cutoff of 115 kya, corresponding with the start of the last glacial period, for 40 sea-level depth intervals, we found that empirical ratios of migrants for 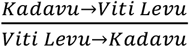 broadly correspond with neutral expectations, suggesting that net migration rates are unlikely to have changed much over the last 115,000 years (Fig. 5). In contrast, empirical migration ratios for 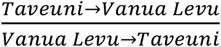 seem consistent with neutral expectations based on present-day sea levels but not expectations averaged over the last 115,000 years (Fig. 5). The empirical net migration ratios for 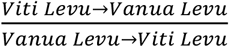 are challenging to interpret given the broad confidence intervals of the empirical data, but the mean empirical migration ratio from the model removing a putatively admixed individual is similar to the value inferred from the centroid-to-centroid distance model averaged over 115,000 years (Fig. 5).

**Figure 5:**
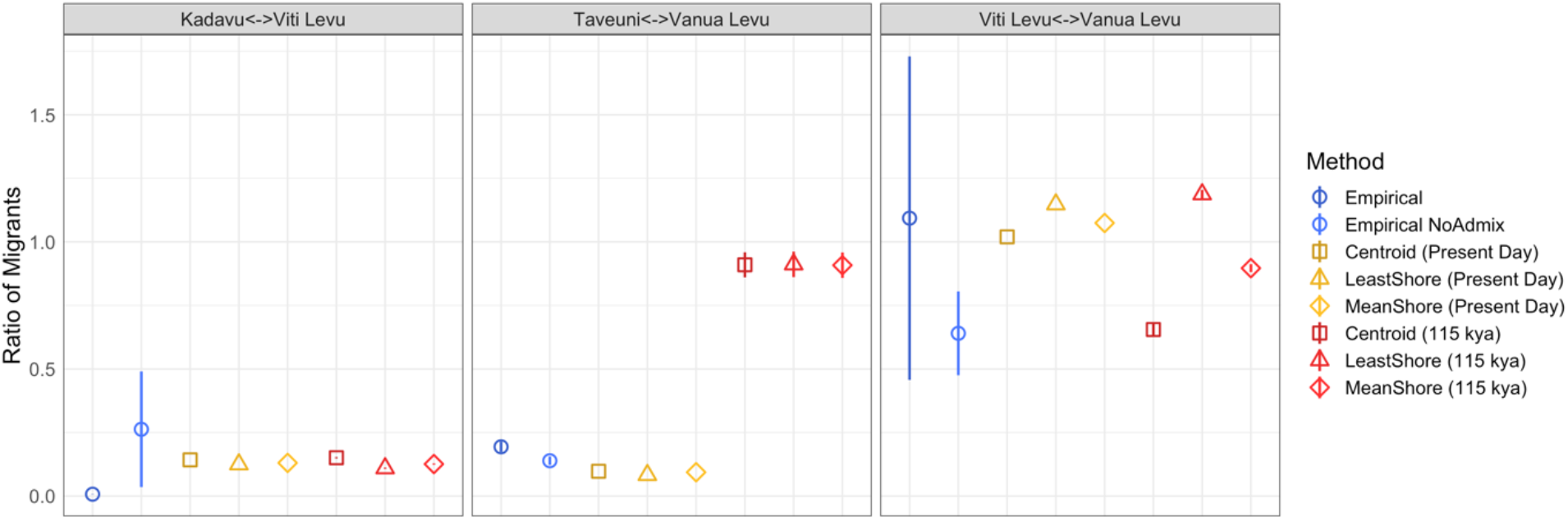
Comparison of empirical estimates of net inter-island migration with contemporary and time-corrected neutral expectations from MacArthur and Wilson (1967), figure adapted from Gyllenhaal et al. 2020.

### 3.4 Spatially explicit forward population genetic simulations

To demonstrate the utility of PleistoDist for spatially explicit simulations, we ran forward-in-time, spatially explicit simulations in SLiM3 (Haller and Messer 2019). We generated raster files for the Samoan archipelago in thousand-year intervals, converted them to .png files using ImageMagick v6.9.7-4, and parameterised spatially explicit Wright-Fisher simulations, allowing individuals to only occupy land exposed during a given time interval. We allowed for two kinds of dispersal: short-distance overland dispersal near the parent and attempted long-distance dispersal (0.2% occurrence probability, using an exponential dispersal kernel, for random dispersal directions). All simulations included spatial competition and mate choice, a metapopulation size of 20,000 individuals, and ran for 50,000 generations. We ran simulations with and without sea-level change, with a generation time of two years. For both dynamic and constant sea-levels, we used eight mean dispersal distances (5, 10, 25, 50, 100, 200, 400, and 800 pixels, ∼4.46 pixels/km), for 100 replicates per sea-level regime/dispersal distance combination. For the last 10 generations of the simulation, we assigned individuals to islands based on their spatial position, and calculated F_ST_ for all island pairs, and per-island nucleotide diversity (π). These 10 generations were averaged to avoid the influence of single migrants on summary statistics. As expected, F_ST_ was negatively correlated with dispersal distance (Fig. 6), and π was positively correlated. The impact of accounting for change in sea-level was most noticeable for islands that were much closer at glacial maxima (i.e Savai’i and Upolu islands, Samoa), but not when dispersal distance was high.

**Figure 6:**
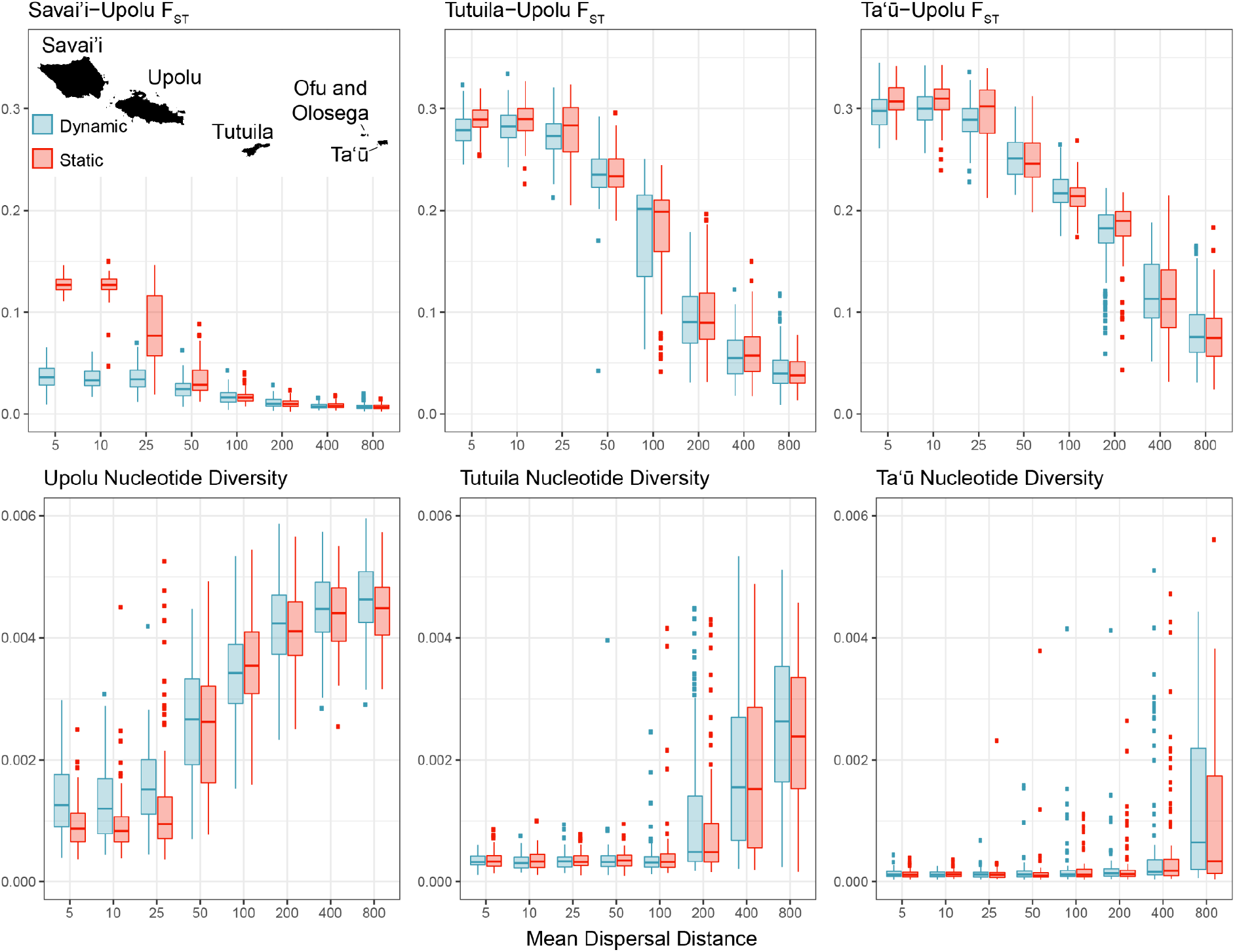
F_ST_ (top row) and π (bottom row) values from spatially explicit simulations across the Samoan Archipelago. Dispersal distance is listed in pixels (∼4.78 pixels/km). Each dispersal distance tested has the results from the dynamic (left, blue) and static (right, red) sea-level models. Inset map has all islands included in the analysis labelled.

To demonstrate the utility of PleistoDist for modelling archipelagic colonisations, we simulated the colonisation of the Solomon Archipelago using the SLiM GUI (Haller and Messer 2019), with initial colonists arriving on Buka/Bougainville islands (Mayr and Diamond 2001). To simulate an expanding colonising population, we used a non-Wright-Fisher simulation, starting with 50 individuals on the easternmost large island in the archipelago. We used a mean long-distance dispersal distance of ∼46.7 km (100 pixels), an exponential long-distance dispersal kernel, and offspring long-distance dispersal probability of 1%. For all simulations (two shown at https://github.com/g33k5p34k/PleistoDistR/tree/main/simulation_scripts), Makira Island was colonised last due to its relative isolation and location at the far eastern end of the archipelago.

### 3.5 Parameterizing migration rates for backwards population genetic simulations

To demonstrate the utility of accounting for migration rate variation between non-connected populations, we used msprime v1.0.0 (Kelleher and Lohse 2020) to model dispersal between Viti Levu and Kadavu islands, Fiji. We used PleistoDist to estimate per interval migration rates using least shore-to-shore distance (Fig. 2) and an exponential dispersal kernel with means of 0 (no dispersal), 10, 20, 40, and 80 km. We calculated population size by multiplying island area for each interval by a density of 10 individuals/km^2^. We either held migration rates and population sizes constant (based on current sea level) or changed them based on sea level in intervals of 5000 years (2500 generations). The initial population split was set to 200,000 generations. At the end of the simulation, we calculated F_ST_ and per-island π foreach sea-level regime. As with the SLiM simulations, the impact of sea level change was more notable when dispersal was low or not present (Fig. 7). These simulations also demonstrate how sea-level change and inter-island dispersal can benefit π on less isolated small island populations (i.e., Kadavu).

**Figure 7:**
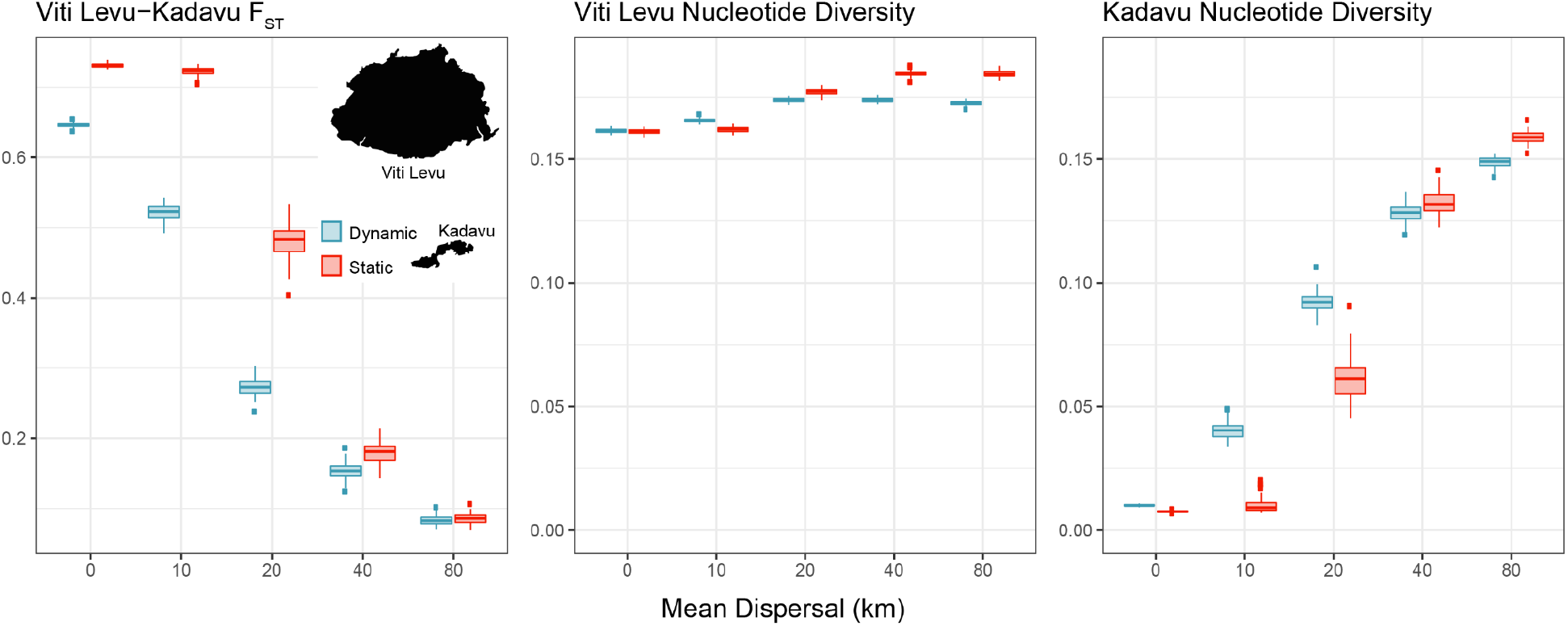
F_ST_ (left panel) and π (middle and right panels) values from msprime simulations for Viti Levu and Kadavu. Each dispersal distance tested has the results from the dynamic (left, blue) and static (right, red) sea-level models.

## 4. Limitations

Despite its broad utility, there are several limitations and assumptions to consider when using PleistoDist. One key assumption is that bathymetry remains constant throughout the timescale of the analysis. This is unlikely to be the case in regions experiencing high levels of tectonic activity. For example, the island of Taveuni in Fiji is likely to have emerged approximately 700 kya (Cronin and Neall 2001), so PleistoDist analyses of Taveuni with a cutoff time close to and exceeding 700 kya are unlikely to be accurate. In addition, PleistoDist is unable to account for the deformation caused by ice sheets, and is thus likely to be less accurate at high latitudes.

## 5. Summary

Although simple in concept, PleistoDist provides users with the ability to calculate island metrics that account for the effects of Pleistocene sea-level change, allowing biogeographers to test multiple hypotheses of evolution and community assembly when coupled with georeferenced empirical data. The flexibility and extensibility of the software, coupled with its numerous possible use-cases, should facilitate the development of more quantitative approaches to the study of island biogeography, and a better understanding of the patterns of species diversification across island archipelagos.

## Data Accessibility

PleistoDist is a free and open-source software package written in R. The source code, example datasets, and a user manual can be found at https://github.com/g33k5p34k/PleistoDistR. The case studies described above are available as vignettes from https://davidbirdtan.com/pleistodist/.

## Authors’ Contributions

DJXT, EFG, and MJA conceived the methodology and concept. DJXT and EFG developed and tested case studies and applications. DJXT developed the package and led the writing of the manuscript, with substantial input from all authors.

